# A programmable dual-targeting di-valent siRNA scaffold supports potent multi-gene modulation in the central nervous system

**DOI:** 10.1101/2023.12.19.572404

**Authors:** Jillian Belgrad, Qi Tang, Sam Hildebrand, Ashley Summers, Ellen Sapp, Dimas Echeverria, Dan O’Reilly, Eric Luu, Brianna Bramato, Sarah Allen, David Cooper, Julia Alterman, Ken Yamada, Neil Aronin, Marian DiFiglia, Anastasia Khvorova

**Affiliations:** RNA Therapeutics Institute, University of Massachusetts Chan Medical School; Worcester, Massachusetts, USA; Department of Neurology, Massachusetts General Hospital; Boston, Massachusetts, USA; Department of Medicine, University of Massachusetts Chan Medical School; Worcester, Massachusetts, USA; Program in Molecular Medicine, University of Massachusetts Chan Medical School; Worcester, Massachusetts, USA

## Abstract

Di-valent short interfering RNA (siRNA) is a promising therapeutic modality that enables sequence-specific modulation of a single target gene in the central nervous system (CNS). To treat complex neurodegenerative disorders, where pathogenesis is driven by multiple genes or pathways, di-valent siRNA must be able to silence multiple target genes simultaneously. Here we present a framework for designing unimolecular “dual-targeting” di-valent siRNAs capable of co-silencing two genes in the CNS. We reconfigured di-valent siRNA – in which two identical, linked siRNAs are made concurrently – to create linear di-valent siRNA – where two siRNAs are made sequentially attached by a covalent linker. This linear configuration, synthesized using commercially available reagents, enables incorporation of two different siRNAs to silence two different targets. We demonstrate that this dual-targeting di-valent siRNA is fully functional in the CNS of mice, supporting at least two months of maximal target silencing. Dual-targeting di-valent siRNA is highly programmable, enabling simultaneous modulation of two different disease-relevant gene pairs (e.g., Huntington’s disease: *MSH3* and *HTT*; Alzheimer’s disease: *APOE* and *JAK1*) with similar potency to a mixture of single-targeting di-valent siRNAs against each gene. This work potentiates CNS modulation of virtually any pair of disease-related targets using a simple unimolecular siRNA.

## INTRODUCTION

Short interfering RNAs (siRNAs) are a powerful therapeutic modality for sequence-specific silencing of target genes. Whereas the siRNA sequence defines the gene target, the chemical architecture of an siRNA dictates stability and delivery *in vivo*. Thus, once an siRNA architecture has been optimized for delivery to a target tissue, any gene with a known sequence in that tissue can, in theory, be targeted by changing the siRNA sequence. Indeed, after establishing an siRNA architecture for delivery to the liver, five siRNA drugs were rapidly developed and FDA-approved to treat liver-related disorders, with many more in clinical trials. Recent advances in siRNA chemistry have led to the development of novel approaches, including lipophilic conjugates and a di-valent scaffold, that enable potent, long-lasting modulation of gene expression throughout the central nervous system (CNS) of both rodent and large animal models (1–4). The di-valent siRNA scaffold, in particular, can induce potent and specific target silencing in the CNS for up to 6 months following direct administration into cerebrospinal fluid (CSF) via a single intracerebroventricular (ICV) injection (1). In the di-valent siRNA scaffold, two asymmetric fully chemically modified siRNA molecules are covalently linked through the 3’ ends of their sense strands. The increased molecular size of di-valent siRNA is a major factor contributing to its prolonged CSF retention and broad distribution throughout the brain (5). However, di-valent siRNA, in its current configuration, can only accommodate two identical siRNA molecules because the arms are grown concurrently during synthesis; and thus, is designed to silence only one gene target.

Single-target modulation by di-valent siRNA may be insufficient for treating neurodegenerative disorders like Huntington’s disease (HD) and Alzheimer’s disease (AD), where pathogenesis is driven by multiple genes or pathways (6, 7). Administering a mixture of different di-valent siRNAs targeting different genes to simultaneously modulate more than one target may represent a viable therapeutic path. However, employing an unimolecular multi-targeting siRNA entity is clinically preferable as it offers a well-defined pharmacokinetic profile upon delivery to the CNS and a more predictable pharmacological profile *in vivo*.

Here, we sought to reconfigure the di-valent scaffold to incorporate different siRNA sequences that, in principle, allow for silencing of two different genes simultaneously. We synthesized dual-targeting di-valent siRNA using an inexpensive, simple linear synthesis scheme where the sense strand arms are grown sequentially during synthesis with an inter-strand linker attaching the arms 5’-3’. We then tested its efficacy in silencing two genes implicated in HD, the disease-causing *HTT* gene and the disease-accelerating *MSH3* gene. HD is caused by a trinucleotide (CAG) repeat expansion in exon 1 of *HTT*, producing toxic *HTT* mRNA and pathogenic HTT protein (8). MSH3, a mismatch repair protein, contributes to further expansion of the inherited CAG expanded region in neurons over a patient’s lifetime (9–12). Increased *HTT* CAG repeat length and rate of expansion lead to earlier disease onset in patients (9, 13, 14) (15, 16), (8, 17). Di-valent siRNAs targeting *Msh3* achieve months-long silencing of Msh3 protein (18), and di-valent siRNAs targeting *Htt* support months-long silencing of Htt protein, in the CNS of mice (1).

We found that dual-targeting di-valent siRNA showed equal efficacy to a mixture of two single-targeting di-valent siRNAs in silencing Msh3 and Htt. When re-programmed to target two AD-associated genes, *Apoe* and *Jak1*, dual-targeting di-valent siRNA achieved efficient silencing for at least 2 months post-injection in the CNS in vivo, similar to that observed in silencing Msh3 and Htt. This study demonstrates the functionality, programmability, and potency of a unimolecular dual-targeting di-valent siRNA scaffold in the CNS, and represents a significant advancement in the design of drug modalities for complex neurodegenerative disorders.

## MATERIALS AND METHODS

### Oligonucleotide synthesis, deprotection, and purification

Oligonucleotides were synthesized by solid-phase phosphoramidite chemistry on a MerMade12 (Biosearch Technologies, Novato, CA) or Dr Oligo 48 (Biolytic, Fremont, CA) using 2’-Fluoro RNA or 2’-*O*-Methyl RNA modified phosphoramidites (ChemGenes, Wilmington, MA, and Hongene Biotech, Union City, CA) (Supplemental Table 1). 5’-(*E*)-Vinyl tetraphosphonate (pivaloyloxymethyl) 2’-O-methyl-uridine 3’-CE phosphoramidite was purchased from Hongene. Phosphoramidites were prepared at 0.1 M in anhydrous acetonitrile (ACN), except for 2’-O-methyl-uridine phosphoramidite dissolved in anhydrous ACN containing 15% dimethylformamide. 5-(Benzylthio)-1H-tetrazole (BTT) was used as the activator at 0.25 M, and coupling time and excess phosphoramidite were 4 mins and 10eq, respectively. All synthesis reagents were purchased from Chemgenes, unless otherwise specified. Unconjugated oligonucleotides were synthesized on 500Å long-chain alkyl amine (LCAA) controlled pore glass (CPG) functionalized with Unylinker terminus. Divalent oligonucleotides (DIO) were synthesized on a custom solid support prepared in-house.

Vinyl-phosphonate-containing oligonucleotides were cleaved and deprotected with 3% diethylamine in ammonium hydroxide, for 20 h at 35℃ with slight agitation. DIO were deprotected with 1:1 ammonium hydroxide and aqueous monomethylamine, for 2 h at 25℃ with slight agitation. The controlled pore glass was subsequently filtered and rinsed with 30 mL of 5% ACN in water and dried overnight by centrifugal vacuum concentration. Purifications were performed on an Agilent 1290 Infinity II HPLC system using Source 15Q anion exchange resin (Cytiva, Marlborough, MA). The loading solution was 20 mM sodium acetate in 10% ACN in water, and elution solution was the loading solution with 1M sodium bromide, using a linear gradient from 30% to 70% in 40 mins at 50°C. Pure fractions were combined and desalted by size-exclusion chromatography with Sephadex G-25 (Cytiva). Purity and identity of fractions and pure oligonucleotides were confirmed by IP-RP LC-MS on an Agilent 6530 Accurate-mass Q-TOF LC-MS.

### Oligonucleotide preparation for in vivo use

Single strands were duplexed by adding a 1:1 ratio of antisense strand to sense strand in water at the desired concentration (10 or 5 nmol di-valent). The antisense and sense mixture was heated at 95°C for 10-20 mins, then removed from heat and allowed to cool at room temperature. A 20% TBE gel (Invitrogen, EC63152BOX) was used to verify successful duplexing. Duplexed siRNAs were dried down using a speed vacuum to remove excess water and then resuspended in calcium and magnesium-enriched artificial CSF (137 mM NaCl, 5 mM KCl, 14 mM CaCl_2_, 2 mM MgCl_2_, 20 mM glucose, 8 mM HEPES, the pH was adjusted to 7.4 with NaOH) to 5 nmol/10 µL or 10 nmol/10 µL, according to the need for individual experiments. Di-siRNA were dosed between (5 nmol) 120 µg to (20 nmol) 440 µg per mouse.

### Animal studies

All animal studies performed were approved by the University of Massachusetts Chan Medical School Institution Animal Care and Use Committee (IACUC PROTO202000010) and used following the National Institutes of Health Guide for the Care and Use of Laboratory Animals. Animals were housed and maintained at pathogen-free animal facilities at UMass Chan Medical School under a 12-hour light/12-hour dark cycle at a controlled temperature (23 ± 1 °C), standard humidity (50% ± 20%), and free access to food and water.

### Stereotactic intracerebroventricular (ICV) injections in mice

FVB/NJ males and females (Jackson Laboratory, Strain #001800) aged 12 weeks (± 2 weeks) were anesthetized with 2.5% isoflurane in oxygen. Their hair at surgical sites were shaved with an electric razor and their heads were placed into the sterile surgical field. A burr was used to drill a small hole in the following coordinates: −0.2 mm posterior, 1.0 mm mediolaterally, the needle was then placed at a depth of −2.5 mm ventrally. A volume of 5 µL was injected per ventricle at a rate of 750 nL/min. Following injection, mice were monitored until sternal.

### Tissue processing

At sacrifice, the brain was harvested and 1.5 x 1.5 mm punches were flash frozen on dry ice or placed in RNALater (935 mL MilliQ water, 700 g Ammonium sulfate, 25 mL 1M Sodium Citrate, 40 mL 0.5 M EDTA, pH 5.2 using H_2_SO_4_) for 24 hours at 4°C, then frozen at −80°C until being processed for subsequent experiments. For mRNA quantification, 1 punch was lysed in 600 µL of QuantiGene assay homogenizing buffer (ThermoFisher Scientific, QS0518) with 0.2 mg/mL proteinase K (ThermoFisher Scientific, AM2546). mRNA was quantified by QuantiGene (ThermoFisher Scientific, QS0016, QS0106) using the manufacture’s protocol. The following QuantiGene Singleplex RNA probes were used from ThermoFisher Scientific: Htt (Murine, SB-14150), Msh3 (Murine, SB-3030208), Hprt (Murine, SB-15463), Jak1 (Murine, SB-3029714), and Apoe (Murine, SB-13611).

### Protein quantification

Protein quantification was performed by western blot analysis. Frozen tissues from striatum or medial cortex were homogenized in 75 µL 10 mM HEPES (pH 7.2) with 250 mM sucrose, 1 mM EDTA + protease inhibitor tablet (Roche #04693116001; complete, EDTA-free) + 1 mM NaF +1 mM Na_3_VO_4_, and sonicated for 10 seconds at 4°C. Bradford assay was used to quantify total protein concentration. 10 µg of protein were first separated by SDS-PAGE and then blotted with the following antibodies: MSH3 (1:500, Santa Cruz, sc-271080), huntingtin (1:2,000, Ab1, aa1-17, (19)), GAPDH (1:10,000, Millipore, AB2302), and β-tubulin (1:5,000, Sigma, T8328). SuperSignal West Pico PLUS chemiluminescence (Pierce) substrate was used to visualize bands. Pixel intensity was manually measured using Image J (Version 1.53t 24, NIH). The average signal intensity was multiplied by the band area to determine the total intensity per band. The band intensity was then normalized to GAPDH or tubulin loading control.

### Guide strand quantification

Peptide nucleic acid (PNA) hybridization assay was used to quantify guide strand accumulation. The PNA probe against the MSH3-targeting antisense strand was ordered from PNA Bio (Thousand Oaks, CA, USA) with the sequence (5’-3’, N-terminal to C-terminal): Alexa546-OO-AAGCAAACTGAAACTGC. Brain lysate (as prepared above) was treated with 3M potassium chloride and pelleted at 4000 ×g for 15 mins to precipitate out Sodium dodecyl sulfate (SDS) from QuantiGene homogenizing buffer (listed above). Alexa546-conjugated PNA probe was incubated with treated lysate, heated to 95°C for 6 mins, and allowed to cool steadily over 30 mins in a thermocycler to anneal the PNA probe to the guide strand of treated tissue. 100 µL per sample of PNA-annealed lysate was then run through an HPLC (Agilent) with a DNAPac PA100 anion-exchange column (ThermoFisher, #043010). Alexa546 fluorescence was recorded, and peaks were integrated to quantify signal. The final guide concentration was quantified by using a standard calibration curve.

### Statistical analysis

All statistics were calculated with GraphPad Prism 10.0.3. Bar graphs are annotated with the sample mean and standard deviation. In vitro experiments were performed in 3 biological replicates. In vivo experiments were performed with n=5-6 mice per group with mRNA and protein experiments performed in technical duplicate or triplicate. One-way ANOVA with Dunnet’s multiple comparisons or with Tukey’s multiple comparisons were used when comparing two or more treatment groups. Two-way ANOVA with Tukey’s multiple comparisons was used when comparing more than two treatment groups across multiple brain regions. P-values are two-sided. When reporting, the following symbols were used: ∗p ≤ 0.05, ∗∗p ≤ 0.01, ∗∗∗p ≤ 0.001, and ∗∗∗∗p ≤ 0.0001.

## RESULTS

### Design unimolecular dual-targeting siRNA scaffold configuration

In the original di-valent siRNA configuration (1), valency was increased using a branched linker for the concurrent synthesis of siRNA sense strands where each coupling reaction was identical for both growing strands (Supplemental Figure 1A). As such, the two siRNAs are linked to each other through the 3’ ends of each sense strand, creating a “branched” configuration (Figure 1A, C, G, Supplemental Figure 1). Chemical strategies can be implemented to adapt this approach to synthesize dual-targeting compounds, but require specialized reagents like reverse phosphoramidites (20) or custom-manufactured linkers with orthogonal protecting groups (21).

**Figure 1:**
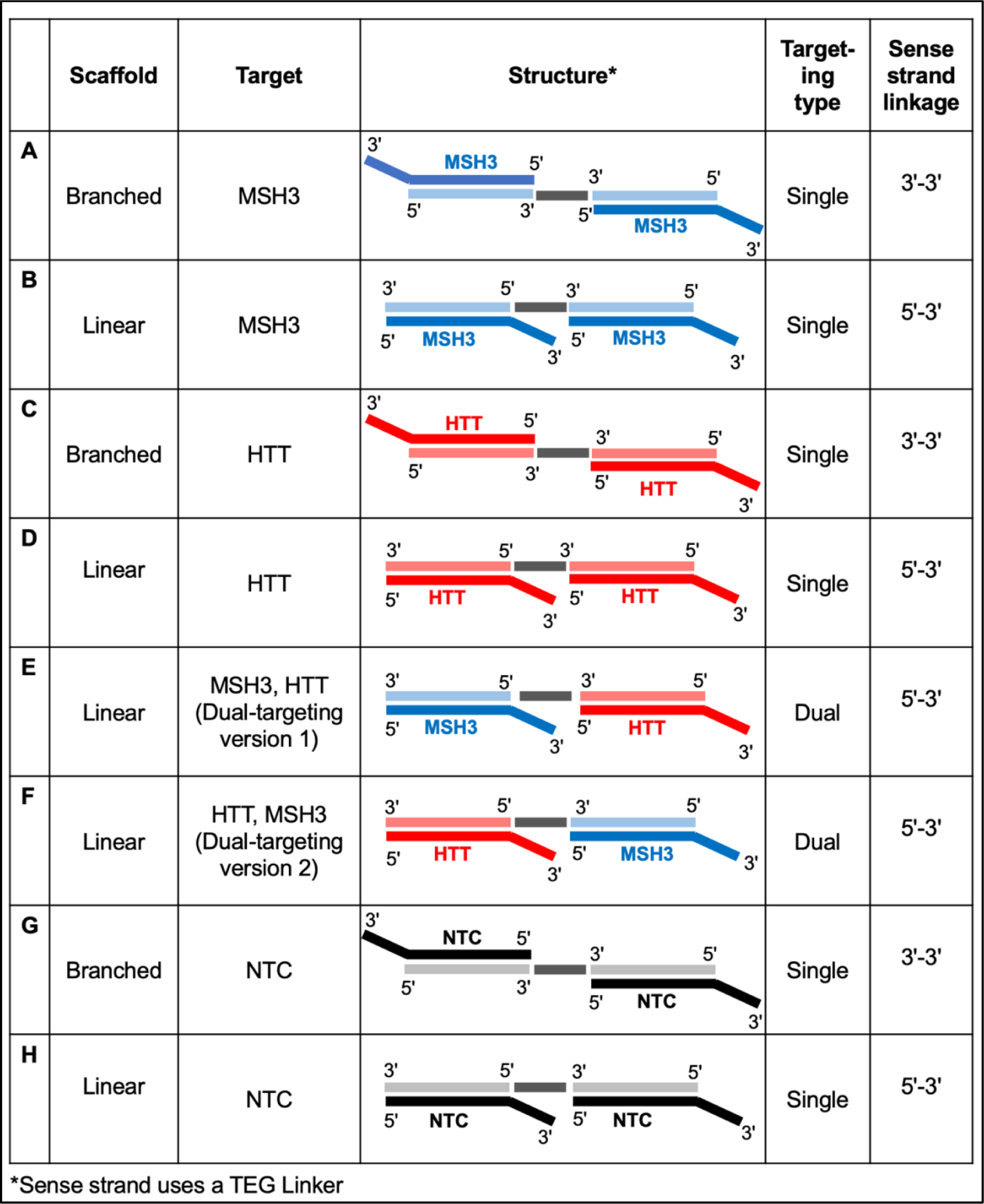
Structures of single- and dual-targeting di-valent siRNA. Branched di-valent siRNA are linked 3’-3’. Linear di-valent siRNA are linked 5’ to 3’. All di-valent siRNA are linked with TEG linker (medium gray bar). Lighter colors represent the sense strand, darker colors represent antisense strand. siRNAs have an antisense strand length of 20 nucleotides and a sense strand length of 16 nucleotides. Sequences and modification patterns are shown in Supplemental Table 1.

To make this approach more accessible and scalable for any research group, we designed a dual-targeting siRNA configuration in which one sense strand is synthesized 3’-5’, a covalent linker is added, and then the second sense strand is synthesized (Supplemental Figure 1B). In this approach, the two sense strand sequences can be unique and are connected 5’-3’, creating a “linear” configuration (Figure 1B, D, E, F, H, Supplemental Figure 1). With this approach, the number of coupling reactions for the single oligonucleotide is doubled, lowering the total yield. However, a sufficient amount of material can be generated to support in vivo studies.

### Linear and branched configurations of di-valent siRNA exhibit comparable efficiency

Branched di-valent siRNA has been shown to achieve sustained (up to 6 months), efficacious silencing of gene targets in the CNS (1, 18, 22, 23). To test the effect of di-valent siRNA configuration on silencing efficacy, we synthesized single-targeting branched and linear configurations against a non-targeting control (NTC) sequence an *Msh3* sequence, or an *Htt* sequence (Figure 1). The di-valent siRNAs were synthesized in-house using previously validated mouse Htt, Msh3, and NTC siRNA sequences (1, 18) in a fully chemically modified pattern for stability against nucleases (24) (Supplemental Table 1).

10 nmol (∼240 μg) of single-targeting branched or linear di-valent siRNAs targeting NTC, Msh3, or Htt were delivered to wild-type (WT) mice via bi-lateral ICV injection (5 nmol per ventricle), resulting in delivery of 20 nmol of active antisense strand to the CNS. One month post-injection, both branched and linear configurations of di-valent siRNA achieved 50-60% target mRNA silencing (Figure 2A,B) and 80-90% target protein silencing (Figure 2C,D) compared to NTC control. There was no significant difference in the level of *Msh3* mRNA (Figure 2A, p=0.7066, two-way ANOVA with Tukey’s multiple comparisons), MSH3 protein (Figure 2C, Supplemental Figure 2, p=0.8229, one-way ANOVA with Tukey’s multiple comparisons), *Htt* mRNA (Figure 2B, p=0.1056, Two-way ANOVA with Tukey’s multiple comparisons), or HTT protein (Figure 2D, Supplemental Figure 2, p=0.9357, one-way ANOVA with Tukey’s multiple comparisons) silencing between the branched and linear configurations. For both *Htt* and *Msh3*, partial mRNA lowering led to near-complete protein silencing, which has been reported previously (1, 23). Nuclear localization of target mRNA is likely responsible for this phenomenon, as siRNA primarily acts in the cytoplasm (25). Using a peptide nucleic acid hybridization assay, we found no difference in antisense strand accumulation between linear and branched di-siRNA in the brain regions probed (Msh3-targeting antisense strand; Figure 2E, Medial cortex: p=0.9953, Striatum: p=0.9836, Thalamus: p=0.9541, Posterior cortex: p=0.9032, two-way ANOVA with Tukey’s multiple comparisons).

**Figure 2:**
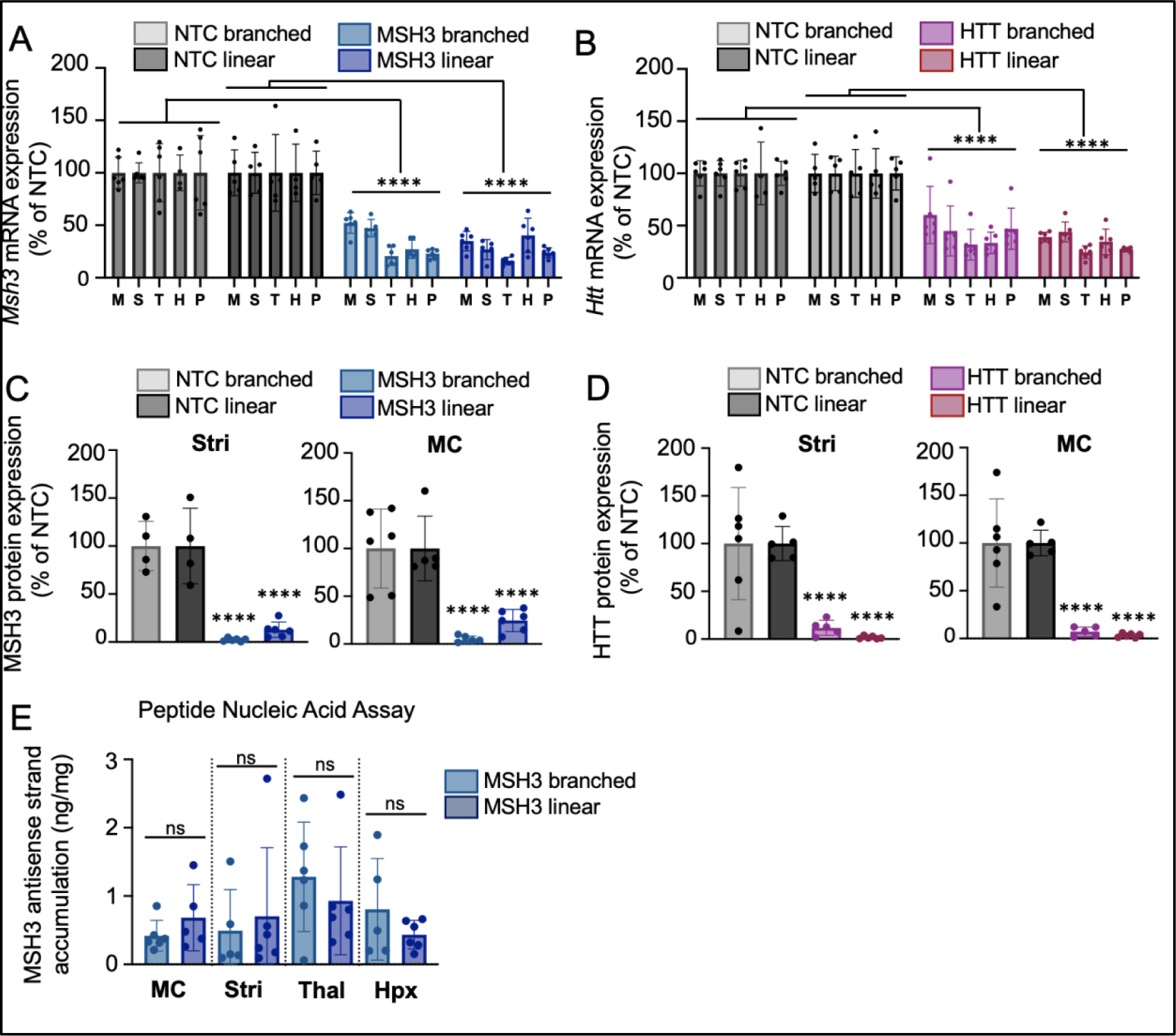
Linear (5’-3’ sense linked) di-valent siRNA performs equivalently to previously validated branched (3’-3’ sense linked) di-valent siRNA. 10 nmol of di-valent siRNA were delivered to wild-type mice via intracerebroventricular injection (5 nmol/5 uL per ventricle). At 1 month post-injection (**A**) *Msh3* mRNA expression was measured by Quantigene assay. (**B**) *Htt* mRNA expression was measured by Quantigene assay. (**C**) MSH3 protein expression was quantified from western blots. (**D**) HTT protein expression was quantified from western blots (**E**) MSH3 antisense strand accumulation was measured by peptide nucleic acid hybridization assay. M/MC-Medial cortex, S/Stri-striatum, T/Thal-thalamus, H/Hpx-hippocampus, P-posterior cortex. Graphs show the mean with standard deviation. Each dot is the average of technical duplicates from a single mouse. N=5-6 mice/treatment group specifically: NTC branched = 6 mice, NTC linear = 5 mice, MSH3 branched = 6 mice, MSH3 linear = 6 mice, HTT branched = 6 mice, HTT linear = 6 mice. NTC is non-targeting di-valent siRNA control. A-B, E: Statistics are two-way ANOVA with Tukey’s multiple comparisons. C-D: Statistics are one-way ANOVA with Tukey’s multiple comparisons. **** indicates p<0.0001.

We next evaluated the safety of linear and branched di-valent siRNA by investigating the impact of treatment on neuroinflammation as measured by GFAP (astrocyte expression) and IBA1 (microglial activation) protein levels. At 1-month post-injection, we observed an increase in striatal IBA1 levels following treatment with branched, but not linear, di-siRNA targeting *Htt* (Supplemental Figure 3A,C; p<0.001, HTT linear vs NTC linear, Ordinary one-way ANOVA with Tukey’s multiple comparisons) which was previously observed and reported to be temporary (1). We also observed a slight but significant increase in cortical IBA1 levels following treatment with linear, but not branched, di-siRNA targeting *Msh3* (Supplemental Figure 3D,F; p<0.01, MSH3 linear vs. NTC linear, Ordinary one-way ANOVA with Tukey’s multiple comparisons). No increase in GFAP was observed for any compound. Taken together, the results suggest configuration of di-valent siRNA had no uniform or consistent impact on neuroinflammation or glia reactivity at 1-month post-injection (Supplemental Figure 3).

Overall, branched and linear di-valent siRNA treatment at a high dose (10 nmol) and short time point (1 month) achieved potent silencing of HTT or MSH3 protein. In the context of this experimental setup, the linear configuration can therefore be used to explore silencing of two different genes simultaneously.

### Dual-targeting di-valent siRNAs are fully functional in the CNS

Using the linear di-valent siRNA configuration, we synthesized a panel of dual-targeting compounds, with one siRNA “arm” targeting Htt and the other siRNA “arm” targeting Msh3. To test whether arm configuration in the dual-targeting siRNA scaffold impacts silencing efficacy, we synthesized two versions of the scaffold where the arm positions of the target sequences were swapped: Msh3 targeting siRNA sequence in arm 1 and Htt targeting sequence in arm 2 (version 1, Figure 1E) and vice versa (version 2, Figure 1F). 10 nmol (∼240 μg) of dual-targeting di-valent siRNA was delivered to WT mice via ICV injection. We delivered 10 nmol linear, single-targeting di-valent siRNAs against NTC only, Htt only, or Msh3 only as controls. At 1 month post-injection, we measured the silencing efficacy of each compound.

Both dual-targeting di-valent siRNA configurations significantly silenced *Msh3* mRNA (>50%) compared to the NTC control (Figure 3A, p<0.0001 NTC linear vs. dual version 1; p<0.0001 NTC linear vs. dual version 2, two-way ANOVA with Tukey’s multiple comparisons) across multiple brain regions. We did observe less potent *Msh3* silencing by dual-targeting di-valent siRNA compared to single-targeting di-valent siRNA (Figure 3A, p=0.0006 MSH3 linear vs. dual version 1, p<0.0001 MSH3 linear vs. dual version 2, two-way ANOVA with Tukey’s multiple comparisons). This difference is likely because the dual-targeting di-valent siRNA delivers half of the amount of Msh3-targeting antisense strand compared to single-targeting di-valent siRNA (10 vs. 20 nmol, respectively).

**Figure 3:**
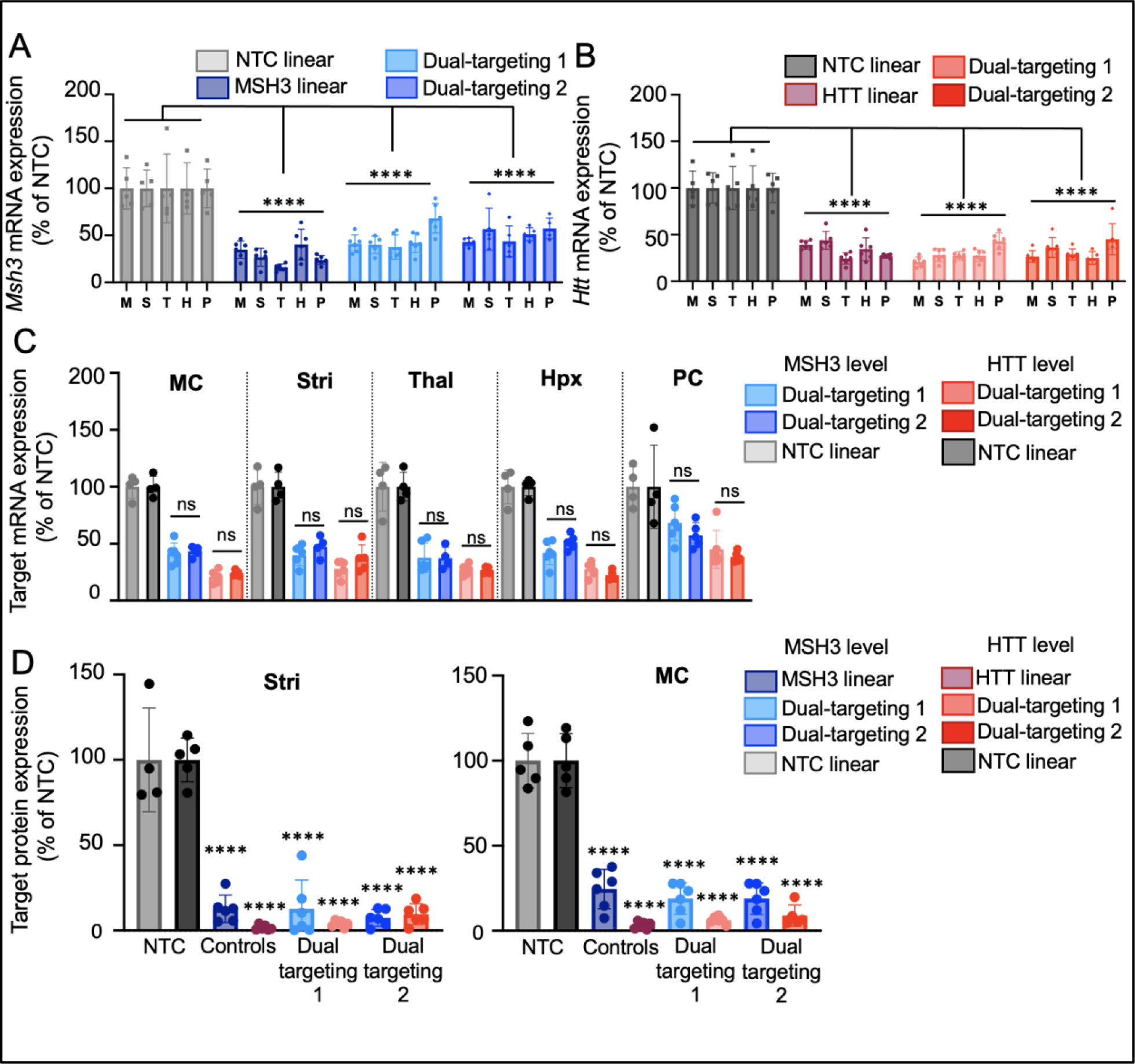
Dual targeting di-valent siRNA reduces mRNA and protein expression of both targets regardless of antisense strand configuration. 10 nmol of di-valent siRNA were delivered to wild-type mice via intracerebroventricular injection (5 nmol/5 uL per ventricle). At 1 month post-injection (**A**) *Msh3* mRNA expression was measured by Quantigene assay. Statistics are two-way ANOVA with Tukey’s multiple comparisons (**B**) *Htt* mRNA expression was measured by Quantigene assay. Statistics are two-way ANOVA with Tukey’s multiple comparisons (**C**) A head-to-head comparison of dual-targeting version 1 and 2. Blue bars are *Msh3* mRNA levels, red bars are *Htt* mRNA levels. (**D**) MSH3 or HTT protein quantified from western blots from the striatum or medial cortex. NTC is a non-targeting control. “Controls” refers to MSH3 or HTT single-targeting di-valent siRNA. Dual-targeting 1 or dual-targeting 2 refers to dual version 1 or 2 respectively. M/MC-Medial cortex, S/Stri-striatum, T/Thal-thalamus, H/Hpx-hippocampus, P-posterior cortex. Sample group size: NTC linear = 5 mice, MSH3 branched = 6 mice, MSH3 linear = 6 mice, HTT branched = 6 mice, HTT linear = 6 mice, dual version 1 = 5 mice, dual version 2 = 6 mice. Graphs are the mean with standard deviation. Each dot is the average of technical duplicates from a single mouse. Ns = no significance, * = p<0.05, ** = p< 0.01, *** = p<0.001, **** = p<0.0001.

Dual-targeting divalent siRNA configurations also significantly silenced *Htt* mRNA versus the NTC control (Figure 3B, p<0.0001, Two-way ANOVA with Tukey’s multiple comparisons), and were not significantly different than single-targeting Htt di-valent siRNA likely due to the potency of the HTT siRNA sequence (Figure 3B, p= 0.4709 HTT linear vs. dual version 1, p=0.7579 HTT linear vs. dual version 2, p=0.9692 dual version 1 vs. dual version 2, two-way ANOVA with Tukey’s multiple comparisons) across multiple brain regions.

Arm configuration had no impact on dual-targeting efficacy. There was no difference in the level of target mRNA silencing when the corresponding antisense strand was in position 1 versus 2 (Figure 3C, D, Supplemental Figure 2, p>0.05 within each brain region dual version 1 vs. dual version 2 for each target, One-way ANOVA with Tukey’s multiple comparisons).

Single- and dual-targeting di-valent siRNA achieved >90% silencing of MSH3 and/or HTT protein compared to the NTC control, with no significant difference in efficacy between configurations for either target (Figure 3C, p>0.9, Two-way ANOVA with Tukey’s multiple comparisons). Once again, there was no difference between the arm position of siRNA within the dual-targeting configuration. These results indicate that, when designing dual-targeting di-valent siRNAs, arm configuration will not impact silencing efficacy.

### Dual-targeting di-valent siRNA exhibit similar gene silencing potency to a mixture of single-targeting di-valent siRNAs

We next sought to compare the silencing efficacies of unimolecular dual-targeting di-valent siRNAs versus a mixture of single-targeting di-valent siRNAs targeting each gene. All siRNAs were synthesized in the linear configuration. Since the dual-targeting arm configuration had no impact on efficacy, we selected dual-targeting version 2 for subsequent experiments. Both the total siRNA dose and the amount of active antisense strand administered may contribute to distribution and efficacy. We, therefore, evaluated all compounds at two different doses, one controlling for the total amount of di-valent siRNA delivered and the other controlling for the total amount of antisense strand (AS) delivered.

We injected a 5 nmol dose (10 nmol total AS strand) or a 10 nmol dose (20 nmol total AS strand) of single-targeting divalent siRNA against Htt only or Msh3 only; a 10 nmol dose (20 nmol total AS strand, 10 nmol each AS strand) or a 20 nmol dose (40 nmol total AS strand, 20 nmol each AS each) of dual-targeting divalent siRNA against Htt and Msh3 or a mixture of single-targeting divalent siRNA targeting Htt only or Msh3 only; or a 20 nmol dose (40 nmol total AS strand) of single-targeting divalent siRNA against NTC only. In this design, we can compare the effect of the total amount of single- and dual-targeting di-valent siRNA and the amount of AS strand per target on silencing efficacy.

Each dose was delivered to WT mice via ICV injection, and target mRNA and protein levels were measured two months post-injection. At both doses, dual-targeting di-valent siRNA silenced *Htt* and *Msh3* mRNA (∼50% silencing) and HTT and MSH3 protein (>90% silencing) compared to NTC control (Figure 4, Striatum, p<0.0001 dual vs. NTC, one-way ANOVA vs. NTC).

**Figure 4:**
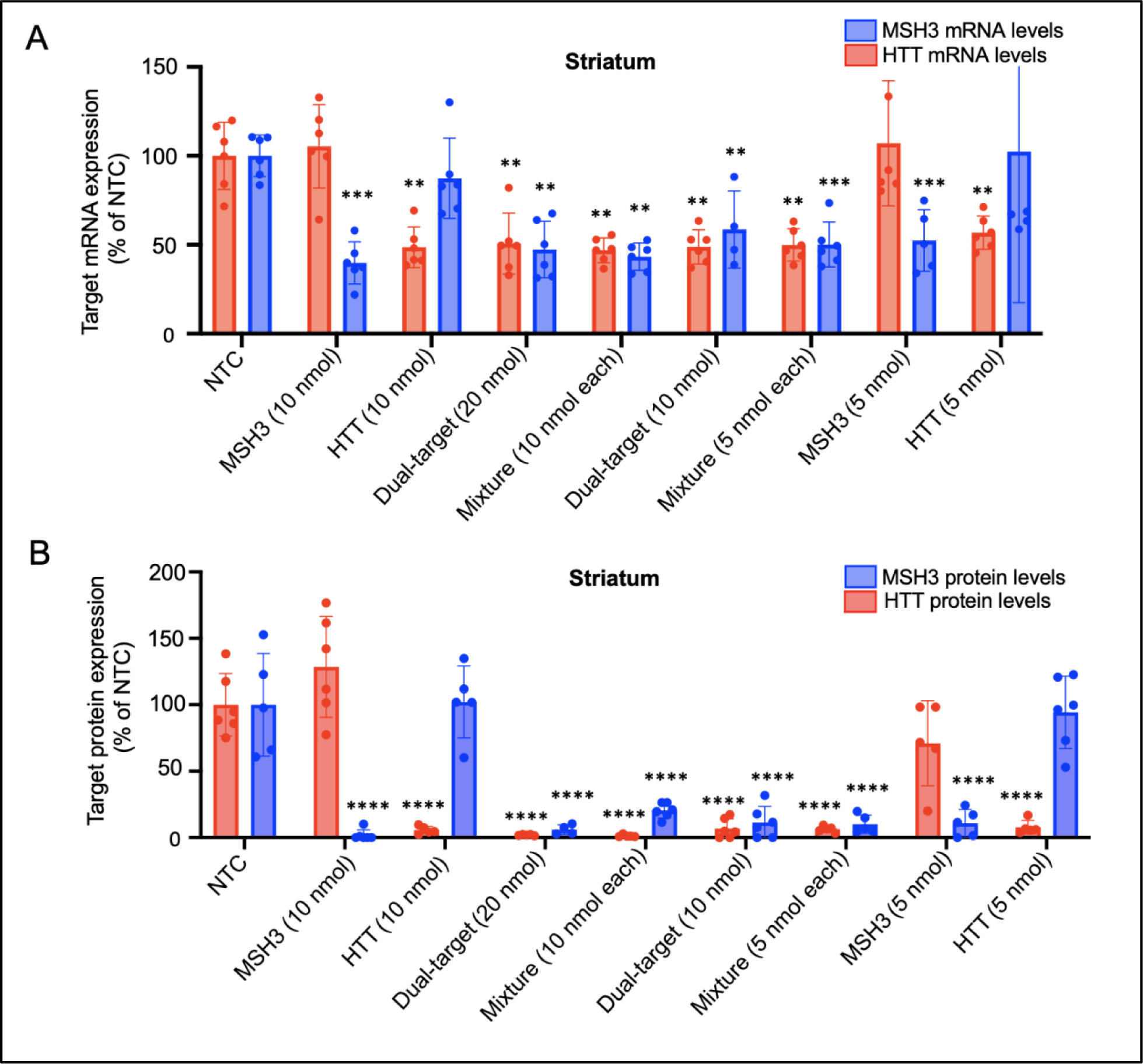
Dual-targeting di-valent siRNA is effective in mouse CNS up to two months in vivo. Di-valent siRNA targeting Msh3 and/or Htt were delivered to wild-type mice via bilateral intracerebroventricular injection. At 2 month post-injection (**A**) Striatal *Htt* (red) or *Msh3* (blue) mRNA expression was measured following treatment with single-targeting, dual-targeting, or a mixture of single-targeting di-valent siRNA. Statistics are one-way ANOVA with Tukey’s multiple comparisons. (**B**) Striatal HTT (red) or MSH3 (blue) protein expression following treatment with single-targeting, dual-targeting, or mixture of single-targeting di-valent siRNA. “NTC” is non-targeting di-valent siRNA control. “MSH3” or “HTT” is single-targeting di-valent control. Dual-target is dual targeting version 2. “Mixture” is a 1:1 combination of Msh3 and Htt single-targeting di-valent siRNAs co-injected as a mixture. Statistics are one-way ANOVA with Tukey’s multiple comparisons. N=5 mice/treatment group. Graphs are the mean with standard deviation. Each dot is the average of technical duplicates from a single mouse. * = p<0.05, ** = p< 0.01, *** = p<0.001, **** = p<0.0001.

At the higher di-valent siRNA dose, single-targeting di-valent siRNA (10 nmol), dual-targeting di-valent siRNA (20 nmol), and the single-targeting di-valent siRNA mixture (20 nmol) each significantly reduced Msh3 mRNA (>50%) and MSH3 protein (>90%) compared to the NTC control (Figure 5A, B, Supplemental Figure 4, p<0.0001 each treatment vs. NTC, Two-way ANOVA with multiple comparisons) and significantly reduced *Htt* mRNA (>50%) and HTT protein (>90%) compared to NTC control (Figure 5C, D, Supplemental Figure 4, p<0.0001 each treatment vs. NTC, Two-way ANOVA with multiple comparisons). There was no significant difference in target mRNA or protein silencing efficacy across any of these treatment groups (p>0.05, two-way ANOVA with Tukey’s multiple comparisons), which all delivered the same amount of target-specific AS strand (20 nmol).

**Figure 5:**
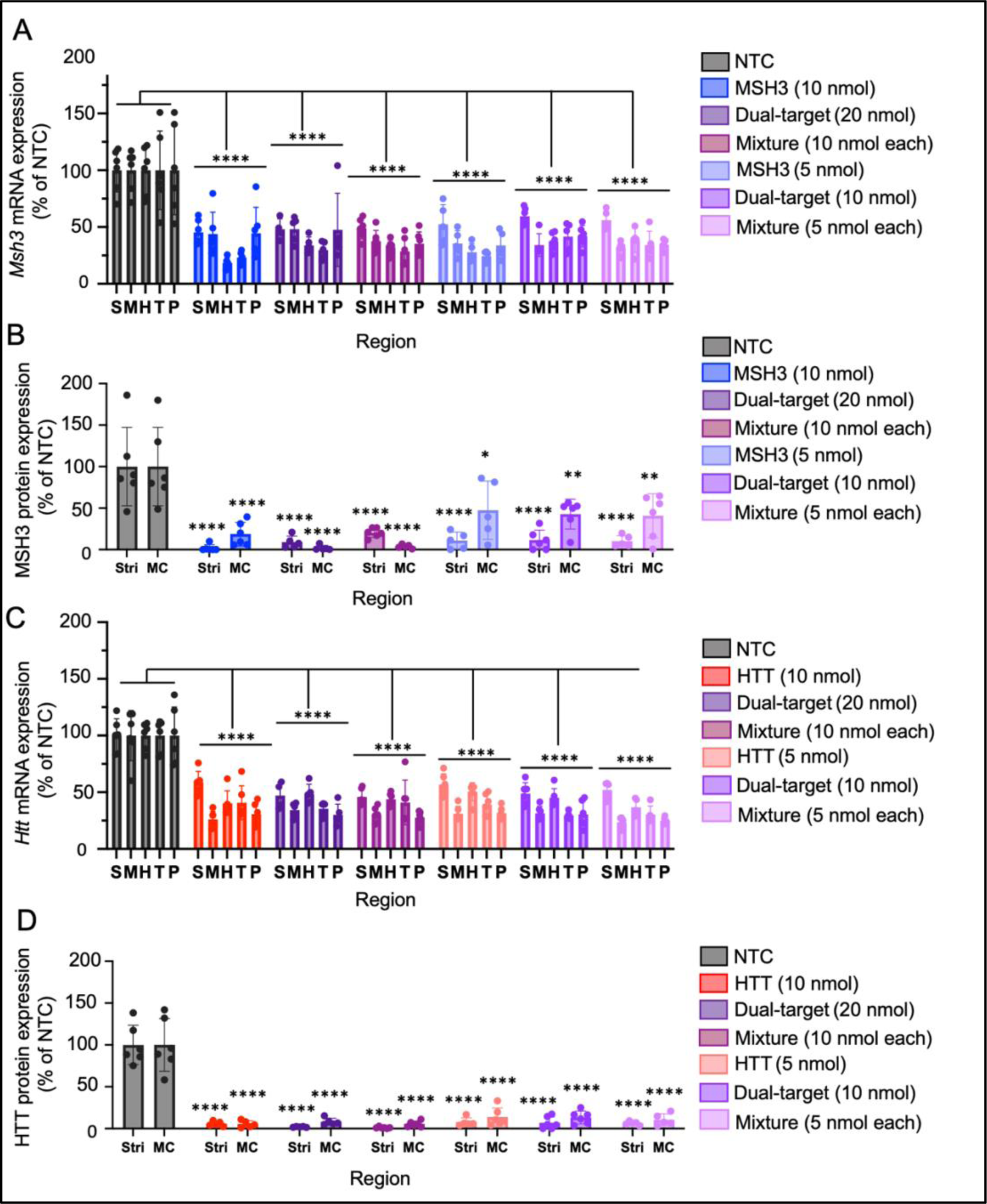
Dual-targeting siRNA works as well as di-valent siRNA mixtures. Di-valent siRNA targeting Msh3 and/or Htt were delivered to wild-type mice via bilateral intracerebroventricular injection. At 2 month post-injection (**A**) *Msh3* mRNA expression was measured by Quantigene assay. Statistics are two-way ANOVA with Tukey’s multiple comparisons (**B**) MSH3 protein expression in the striatum and medial cortex was quantified from western blot. Statistics are two-way ANOVA with Tukey’s multiple comparisons (**C**) *Htt* mRNA expression was measured by Quantigene assay. Statistics are two-way ANOVA with Tukey’s multiple comparisons (**D**) HTT protein expression in the striatum and medial cortex was quantified from western blot. Statistics are two-way ANOVA with Tukey’s multiple comparisons. “NTC” is non-targeting di-valent siRNA control. “MSH3” or “HTT” is single-targeting di-valent control. Dual-target is dual targeting version 2. “Mixture” is a 1:1 combination of Msh3 and Htt single-targeting di-valent siRNAs co-injected as a mixture. M/MC-Medial cortex, S/Stri-striatum, T/Thal-thalamus, H/Hpx-hippocampus, P-posterior cortex. N=5 mice/treatment group. Graphs are the mean with standard deviation. Each dot is the average of technical duplicates from a single mouse. * = p<0.05, ** = p< 0.01, *** = p<0.001, **** = p<0.0001.

At the lower di-valent siRNA doses, single-targeting di-valent siRNA (5 nmol), dual-targeting di-valent siRNA (10 nmol), and the single-targeting di-valent siRNA mixture (10 nmol) each significantly reduced *Msh3* mRNA (>50%) and MSH3 protein (50-90%) compared to NTC control (Figure 5A, B, Two-way ANOVA with multiple comparisons) and significantly reduced *Htt* mRNA (>50%) and HTT protein (>90%) compared to NTC control (Figure 5C, D, p<0.0001 each treatment vs. NTC, Two-way ANOVA with multiple comparisons). There was no significant difference in target mRNA or protein silencing efficacy across treatment groups (p>0.05, two-way ANOVA with Tukey’s multiple comparisons), which all delivered the same amount of target-specific AS strand (10 nmol).

Comparing the high and low doses within each configuration, we observed a reduction in the level of the MSH3, but not the HTT, protein silencing (∼50% vs 80%) in the medial cortex, similar to what has been previously reported (18).

For all treatment groups that delivered a 10 nmol di-valent siRNA dose (with varying AS strand amounts), there was no difference in the level of *Msh3* mRNA silencing between dual-targeting divalent siRNA, single-targeting divalent siRNA against Msh3, and the single-targeting di-valent siRNA mixture against Msh3 and Htt (Figure 5A, p>0.05, two-way ANOVA with multiple comparisons). At the protein level, however, dual-targeting divalent siRNA and the single-targeting di-valent siRNA mixture did perform worse than single-targeting divalent siRNA (Figure 5B, Supplemental Figure 4) in the medial cortex. This is likely because the Msh3-targeting sequence is less potent; and thus, the lower AS strand amount in the dual-targeting divalent siRNA and the single-targeting di-valent siRNA mixture (compared to the single-targeting divalent siRNA) contributed to less potent silencing at the protein level.

For all treatment groups that delivered a 10 nmol di-valent siRNA dose (with varying AS strand amounts), there was no difference in the level of *Htt* mRNA and HTT protein silencing between dual-targeting divalent siRNA, single-targeting divalent siRNA against Htt, and single-targeting di-valent siRNA mixture against Msh3 and Htt (Figure 5C,D; Supplemental Figure 4). Each group also exhibited significant Htt silencing compared to NTC. Here, the HTT-targeting sequence is potent and active across a wide range of doses; and thus, dual-targeting divalent siRNA and the single-targeting di-valent siRNA mixture can maintain maximum efficacy even with a lower AS strand amount (compared to the single-targeting divalent siRNA).

Overall, this study demonstrates that increasing the total amount of siRNA delivered while keeping the AS strand dose the same (e.g., 10 nmol Htt-targeting AS strand in a 10 nmol siRNA dose vs. 10 nmol Htt-targeting AS strand in a 20 nmol siRNA dose) did not enhance target mRNA or protein silencing at the doses and time points tested. As long as the delivered AS strand doses were identical, the knockdown efficacy was the same.

### Dual-targeting di-valent siRNA configurations can be applied to multiple target sequences

To test whether the dual-targeting divalent siRNA scaffold could be reprogrammed to silence other disease-relevant genes, dual-targeting divalent siRNAs and controls were synthesized using two previously validated siRNA sequences, Apoe and Jak1 (Figure 6A) (22, 26). Apolipoprotein E (APOE) is a well-characterized, leading risk factor for AD (27), and Janus kinase 1 (JAK1) is a major signal transduction mediator of many inflammatory pathways, including the interferon pathway (26, 28). Both APOE and JAK1 are promising therapeutic targets for AD (29–32), which is associated with harmful neuroinflammation (33–36). To test the efficacy of dual-targeting divalent siRNA against ApoE and Jak1 in silencing the expression of both genes, we delivered 20 nmol dual-targeting di-valent siRNA to WT mice via ICV injection. Single-target divalent siRNA targeting ApoE only (20 nmol) or NTC only (20 nmol) were used as controls.

**Figure 6:**
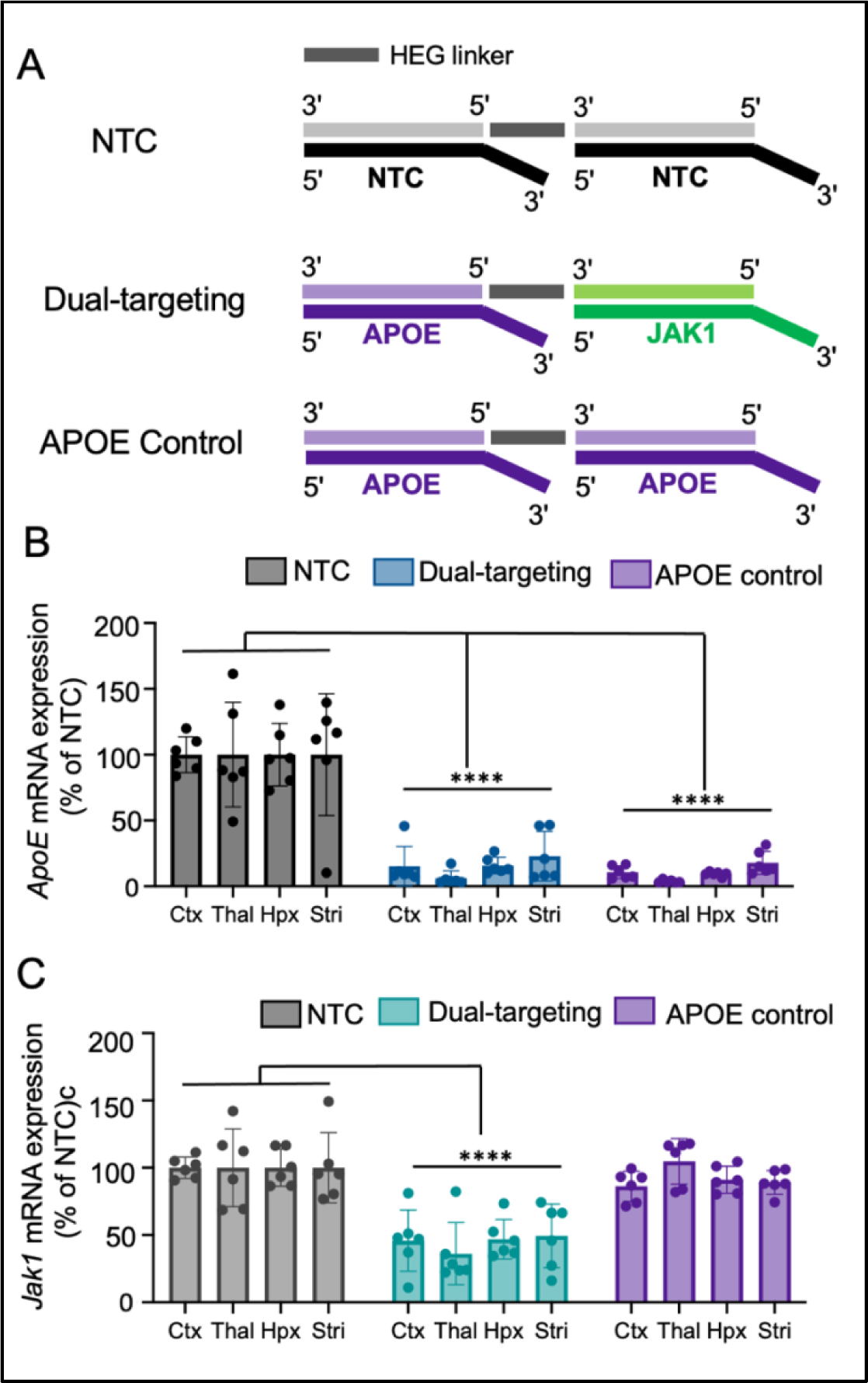
Dual-targeting siRNAs are programable and functional across sequences in vivo. (**A**) 10 nmol of di-valent siRNA were delivered to wild-type mice via intracerebroventricular injection (5 nmol/5 uL per ventricle). NTC, dual-targeting, or APOE control structures used for the experiment. Sequences and modification patterns are shown in Supplemental Table 1. At 2 month post-injection (**B**) *ApoE* mRNA expression was quantified by QuantiGene assay. Statistics are two-way ANOVA with Tukey’s multiple comparisons (**C**) *Jak1* mRNA expression was quantified by QuantiGene. Statistics are two-way ANOVA with Tukey’s multiple comparisons. NTC is non-targeting di-valent siRNA control. N=5 mice/treatment group. Graphs are the mean with standard deviation. Each dot is the average of technical duplicates from a single mouse. * = p<0.05, ** = p< 0.01, *** = p<0.001, **** = p<0.0001.

At two months post-injection, dual-targeting and single-targeting divalent siRNA both induced >80% *Apoe* mRNA silencing compared to NTC control (Figure 6B, p<0.001, Two-way ANOVA with Tukey, multiple comparisons), with no significant difference between the two treatments (Figure 6B, p=0.7292 dual vs. APOE, Two-way ANOVA with Tukey’s multiple comparisons). Dual-targeting divalent siRNA, but not single-targeting divalent siRNA against Apoe, induced >50% *Jak1* mRNA silencing compared to NTC control (Figure 6C, p<0.0001, NTC vs. dual, p=0.3811 APOE vs NTC, Two-way ANOVA with Tukey’s multiple comparisons). Taken together, the technology reported here can be applied to a wide range of target combinations.

## DISCUSSION

Here we characterize the first in vivo ability of multi-targeting siRNA in the CNS. Single gene targeting siRNA including di-valent (1) and lipid-conjugated (2, 4) approaches have transformed the development of nucleic acid therapeutics in neurological diseases. Still, in complex neurodegenerative diseases, single-gene targeting may be insufficient. Further, multi-targeting has been studied extensively systemically (37) but has not yet been successful in the CNS. To address the need for a multi-targeting scaffold in the CNS, we used a linear synthesis scheme for di-valent siRNA. In this linear synthesis, di-valent siRNA sense strands are grown sequentially, rather than concurrently, and attached through a linker to create a dual-targeting di-valent siRNA where each arm has a unique sequence. We show that both linear and branched configurations of di-valent siRNAs are fully functional in vivo, potently modulating the expression of two genes involved in HD (i.e., *HTT* and *MSH3*) and two genes involved in AD (i.e., *APOE* and *JAK1*). The level of silencing by dual-targeting di-valent siRNA was comparable to that of single-targeting di-valent siRNAs administered individually or as a mixture. Our findings establish a new therapeutic tool for multi-target modulation in the CNS.

Several chemical strategies can generate dual-targeting di-valent siRNA, but most require the use of specialized reagents like reverse phosphoramidites and orthogonal protection groups. Here we synthesized a linear configuration of di-valent siRNA (the two sense strands are covalently attached by an inter-strand linker through a linear 3’ to 5’ stepwise synthesis) using commercially available phosphoramidites readily accessible to a wider range of laboratories. At a high dose (10 nmol di-valent) and over a short time frame (1 month), linear di-valent siRNA was as active as the branched di-valent siRNA (concurrent synthesis of identical siRNA arms; sense strands linked 3’-3’). Future work should compare the efficacy of each configuration at different doses and longer durations.

We also showed that the nature (i.e., MSH3, HTT, APOE, and JAK1) and positioning (arm 1 vs. arm 2) of the siRNA sequences in the linear dual-targeting di-valent siRNA scaffold do not impact maximal silencing efficacy. Thus, this technology is highly programmable, making it applicable to many complex neurological disorders.

A major advantage of using unimolecular dual-targeting di-valent siRNA over a mixture of two single-targeting di-valent siRNAs is that safety, efficacy, durability, and off-target validations may only be needed for one therapeutic entity instead of two. This efficiency could quicken the progression of compounds toward clinical trials in patients. Using a unimolecular entity also results in uniform CNS distribution and uptake of both siRNA molecules to the same target cells, ensuring both biological pathways are targeted in a 1:1 ratio in each cell. This uniformity is crucial for diseases like HD, where inhibition of both somatic expansion and HTT expression is needed in each cell to disrupt existing toxic HTT species and reduce future production of these species. Nevertheless, there may be certain disease contexts where the desired target knockdown stoichiometry may not be 1:1. Based on the timeline and higher doses presented in this study, the dual-targeting di-valent siRNA scaffold does produce maximum silencing of both targets, but the use of a lower dose or a less potent sequence or mismatches, could enable target-knockdown ratios other than 1:1. Indeed, in the present study, the Msh3-targeting sequence was slightly less potent than HTT. In the lower dose group, this translated to Htt mRNA and protein, but not Msh3 protein, in the medial cortex remaining silenced at two months post-injection. Titrating sequences in a single-targeting di-valent siRNA mixture could also be used to achieve different knockdown ratios. The work presented here demonstrates a proof-of-concept that can now be applied, optimized, and adapted to tailor dual-targeting therapeutics to the unique needs of different neurological diseases.

Controlling for multi-targeting experiments is complex. Both bulk siRNA dose and active AS dose need to be considered. In general, we found increasing per-target AS amounts, rather than total amount of siRNA delivered, increased the potency of each individual siRNA sequence. The observed effects of AS dose and bulk dose on silencing efficacy may be different in the context of other sequences or when considering systemic rather than CNS delivery, and should be tested in future work.

Traditionally, oligonucleotide therapeutics have been applied to disorders where a single gene causes the disease (38). However, it has become increasingly clear that, for some monogenetic disorders, pathogenesis or progression is associated with multiple pathways (13). It is now understood that effectively treating HD might require targeting both the disease-causing HTT gene and disease-accelerating genes (e.g., the mismatch repair pathway) (13, 39, 40).

Furthermore, modulating multiple genes in the mismatch repair pathway (i.e., PMS1, MLH1, MLH3, MSH3, etc.) might be necessary to block or reverse *HTT* expansion or to target different symptoms (39, 41–43). The di-valent siRNAs developed in this work provide an avenue to develop more complex multi-targeting siRNAs, including tri- and tetra-valent compounds. The size, delivery mechanisms, and structure considerations of these multi-targeting siRNAs must be further explored in future work.

The concept of multi-targeting systemically has been considered previously. Alnylam Pharmaceuticals and others have demonstrated that two siRNAs (Patent US9187746B2) can be covalently linked together and conjugated to a single GalNAc moiety (which targets siRNAs to liver) to silence two genes. In this context, the oligonucleotide to conjugate ratio differs between the dual-targeting (2:1) and single-targeting (1:1) configurations. This ratio change in dual-targeting GalNAc-conjugated siRNAs can impact distribution and lead to slightly lower potency. These differences might be even more pronounced in the context of siRNAs conjugated to lipid moieties, where the ratio between lipophilic conjugate and charged RNA is known to alter distribution (2, 44–47). Unlike GalNAc-or lipid-conjugated siRNAs, di-valent siRNA uniquely lends itself to multi-targeting. Single- and dual-targeting di-valent siRNAs have identical size, charge, and chemical modifications, only differing in their target sequence (1, 24). Since the chemistry and structure of the siRNA (not sequence) are the main factors driving distribution (24), the performance of these compounds in vivo is expected to be consistent. The present study demonstrates this in mice one to two months post-injection. Expanding these observations to large animal models is necessary.

In conclusion, the field’s capacity to use RNA interference-based therapeutics is maturing, with a growing consideration for applying these technologies to complex diseases. Given the biological complexity of neurodegenerative conditions, modulation of multiple pathways will likely be required for the development of disease-modifying drugs. This work unveils the fundamental design principles and introduces an accessible chemical framework for developing therapeutic siRNAs to enable multi-target modulation in the CNS.

## SUPPLEMENTARY DATA STATEMENT

Supplementary Data are available at NAR online.

## DATA AVAILABILITY

The data underlying this article is available upon request to the corresponding author.

