## Supplemental Data for "A programmable dual-targeting di-valent siRNA scaffold supports potent multi-gene modulation in the central nervous system"

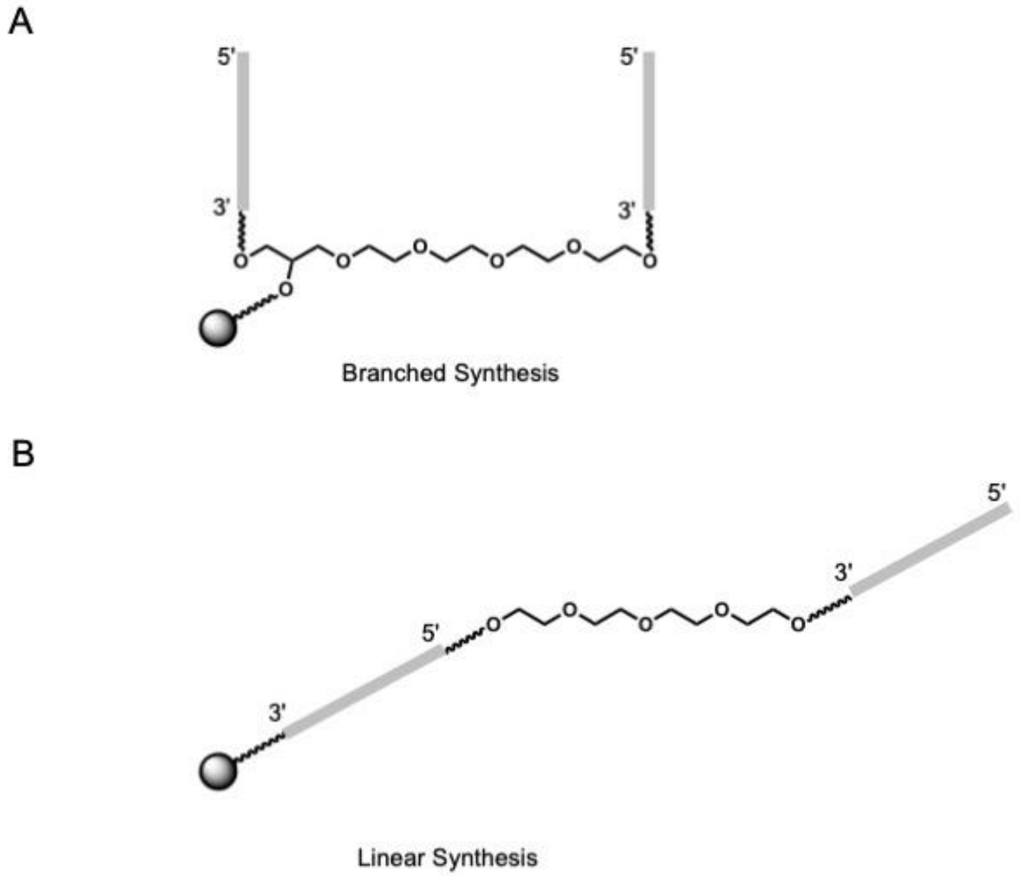


**Supplemental Figure 1: Schematic for synthesis of branched and linear configurations of di-valent siRNA.** (A) Solid phase synthesis schematic for branched-configuration di-valent siRNA. (B) Solid phase synthesis schematic for linear-configuration di-valent siRNA. The gray ball represents the solid phase support and the gray bars are the growing oligonucleotide strands.

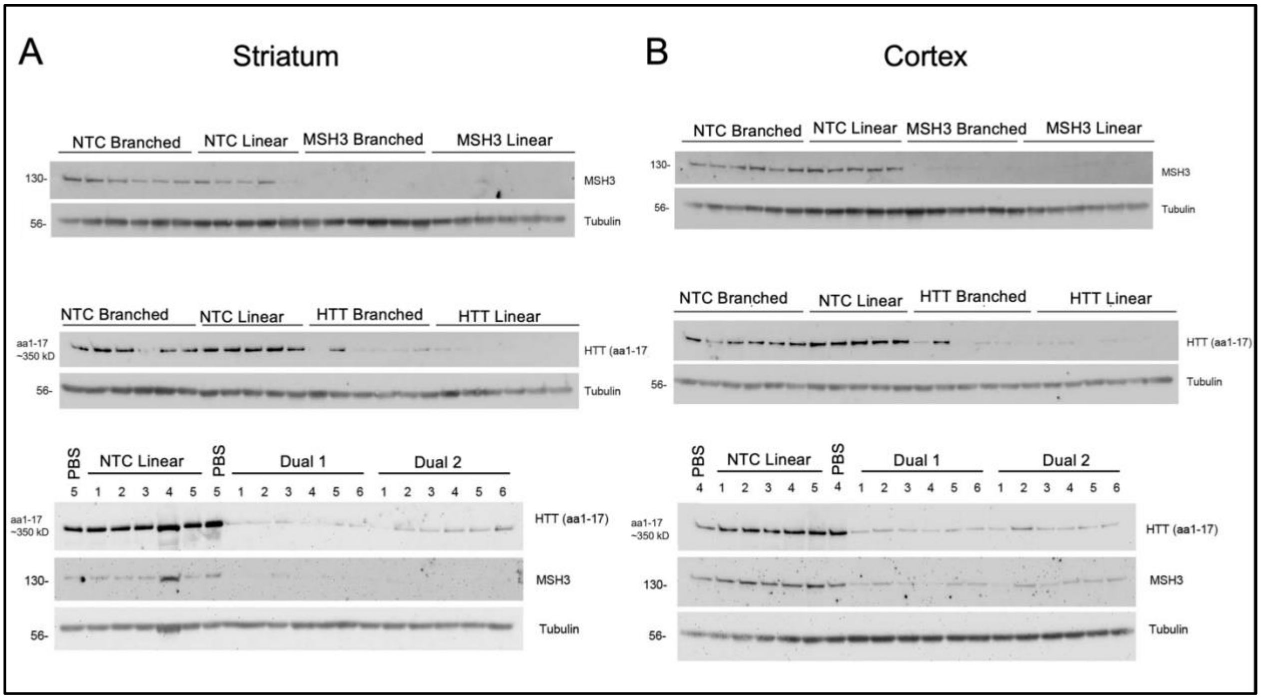


**Supplemental Figure 2: Dual targeting siRNA induces potent silencing of both targets one-month post-injection in vivo** (A) Raw western blots from (A) Striatum and (B) Medial cortex of wild type mice one-month post bilateral intracerebroventricular injection of 10 nmol di-valent siRNA treatments. Samples were blotted with anti-HTT or anti-MSH3 antibodies with tubulin used as a loading control. Numbers along the top indicate individual mouse samples.

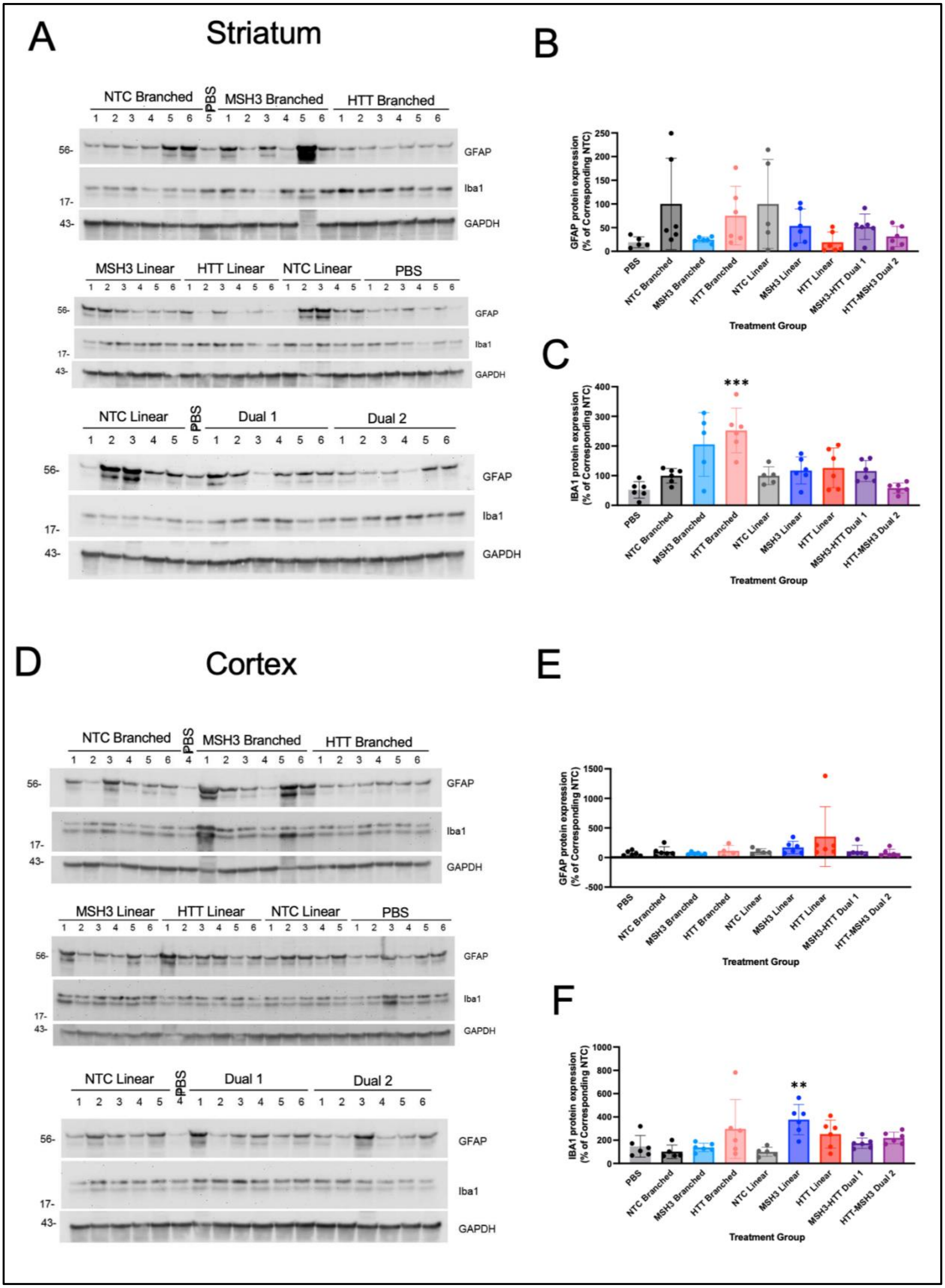
A
Striatum
NTC Branched
PBS
MSH3 Branched
HTT Branched
GFAP
Iba1
GAPDH
MSH3 Linear
HTT Linear
NTC Linear
PBS
Dual 1
Dual 2
B
GFAP protein expression (% of Corresponding NTC)
Treatment Group
C
IBA1 protein expression (% of Corresponding NTC)
***
D
Cortex
E
F
**
PBS
NTC Branched
MSH3 Branched
HTT Branched
NTC Linear
MSH3 Linear
HTT Linear
MSH3-HTT Dual 1
HTT-MSH3 Dual 2

**Supplemental Figure 3: No increase in neuroinflammation occurs with dual-targeting linear di-valent siRNA.** (**A**) Raw western blots form mouse stiatum probing against GFAP or IBA1 protein in wild type mice one-month post bilateral intracerebroventricular injection of 10 nmol di-valent siRNA treatment groups. (**B**) Quantified GFAP protein in the striatum per treatment group. Values are GFAP signal, normalized to tubulin loading control normalized to NTC treated groups. (**C**) Quantified IBA1 protein per treatment group. Values are IBA1 signal, normalized to tubulin loading control normalized to NTC treated groups. (**D**) Raw western blots from mouse cortex probing against GFAP or IBA1 protein in wild-type mice one-month post bilateral intracerebroventricular injection of 10 nmol di-valent siRNA treatment group. (**B**) Quantified GFAP protein in the cortex per treatment group. Values are GFAP signal, normalized to tubulin loading control normalized to NTC treated groups. (**C**) Quantified cortical IBA1 protein per treatment group in the cortex. Values are IBA1 signal, normalized to tubulin loading control normalized to NTC treated groups (siRNA$^{NTC/NTC}$ or siRNA$^{NTC–NTC}$). Numbers along the top indicate individual mouse samples. Statistics are two-way ANOVA with Tukey's multiple comparisons, ** = $p< 0.01$, *** = $p<0.001$

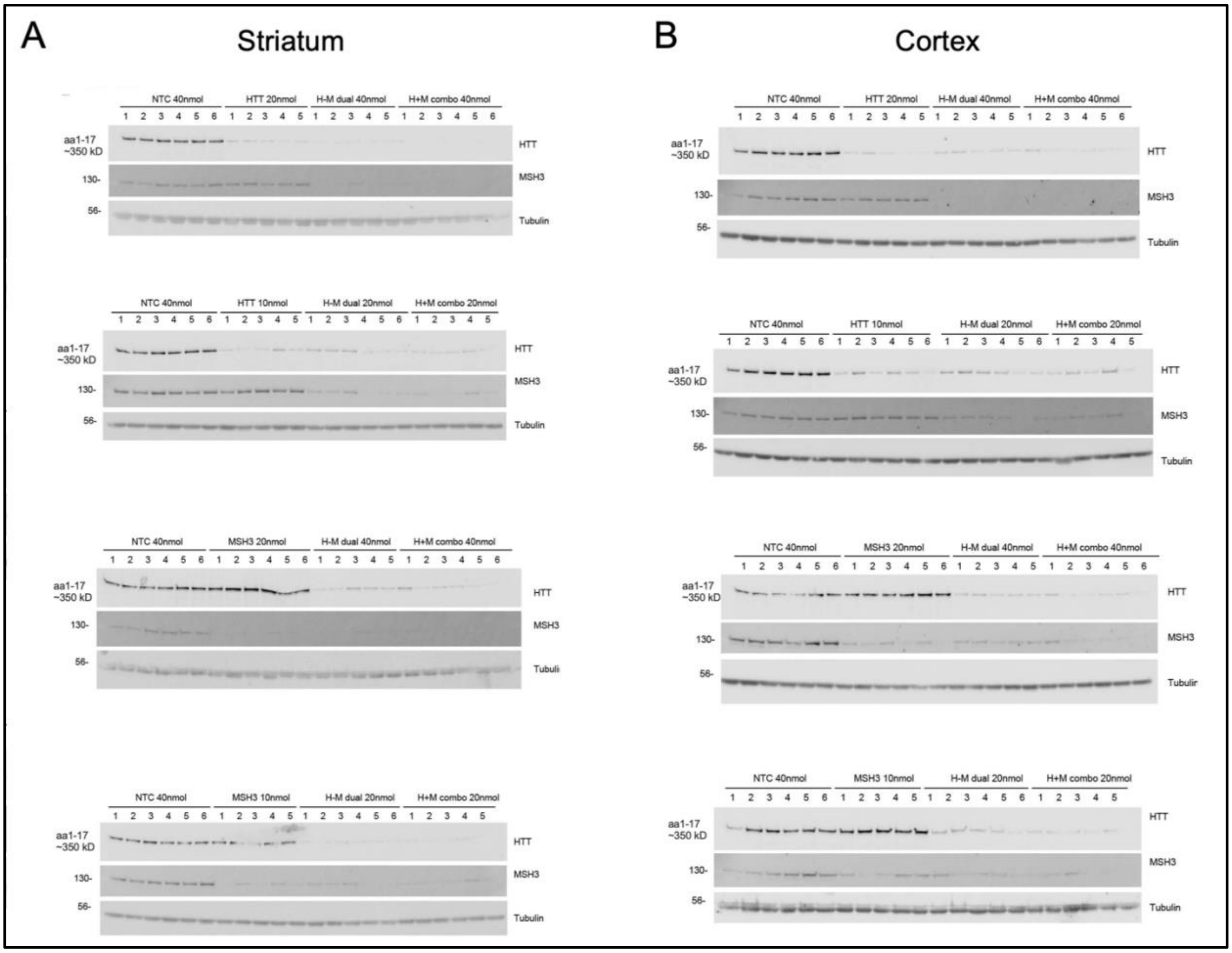


**Supplemental Figure 4: Dual targeting di-valent siRNAs are functional at least two months in vivo and works as well as di-valent siRNA mixture.** Samples are from wild-type mouse two-months post bilateral injection with respective treatment groups. Raw western blots from (A) Striatum or (B) Medial cortex HTT or MSH3 with tubulin loading control. Numbers along the top indicate individual mouse samples.

| Chemical Modification Patterns and Sequences of siRNA | | |
|---|---|---|
| **Oligo Name** | **Strand Name** | **Sequences** |
| NTC Branched | NTC_AS_P3_VP_21mer_3'mf | V(mU)#(fA)#(mA)(fU)(fC)(fG)(mU)(fA)(mU)(fU)(mU)(fG)(mU)(fC)#(mA)#(fA)#(mU)#(mC)#(mA)#(mU)#(fU) |
| | NTC_S_P3_16mer_DIO | (mU)#(mU)#(mG)(fA)(mC)(fA)(mA)(fA)(mU)(fA)(mC)(mG)(mA)(fU)#(mU)#(mA)-DIO |
| NTC Linear | NTC_AS_P3_VP_21mer_3'mf | V(mU)#(fA)#(mA)(fU)(fC)(fG)(mU)(fA)(mU)(fU)(mU)(fG)(mU)(fC)#(mA)#(fA)#(mU)#(mC)#(mA)#(mU)#(fU) |
| | NTC_S_P3_16mer_DIOTEG_5'-3' | (mU)#(mU)#(mG)(fA)(mC)(fA)(mA)(fA)(mU)(fA)(mC)(mG)(mA)(fU)#(mU)#(mA)(TEG)(mU)#(mU)#(mG)(fA)(mC)(fA)(mA)(fA)(mU)(fA)(mC)(mG)(mA)(fU)#(mU)#(mA) |
| MSH3 Branched | MSH3_1000_P3_as_21mer_3'mf | V(mU)#(fG)#(mC)(fA)(fG)(fU)(mU)(fU)(mC)(fA)(mG)(fU)(mU)(fU)#(mG)#(fC)#(mU)#(mU)#(mC)#(mA)#(fU) |
| | MSH3_1000_P3_s_16mer_DIO | (mG)#(mC)#(mA)(fA)(mA)(fC)(mU)(fG)(mA)(fA)(mA)(mC)(mU)(fG)#(mC)#(mA)-DIO |
| MSH3 Linear | MSH3_1000_P3_as_21mer_3'mf | V(mU)#(fG)#(mC)(fA)(fG)(fU)(mU)(fU)(mC)(fA)(mG)(fU)(mU)(fU)#(mG)#(fC)#(mU)#(mU)#(mC)#(mA)#(fU) |
| | MSH3_1000_P3_s_16mer_DIOTEG_5'-3' | (mG)#(mC)#(mA)(fA)(mA)(fC)(mU)(fG)(mA)(fA)(mA)(mC)(mU)(fG)#(mC)#(mA)(TEG)(mG)#(mC)#(mA)(fA)(mA)(fC)(mU)(fG)(mA)(fA)(mA)(mC)(mU)(fG)#(mC)#(mA) |
| HTT Branched | HTT_AS_P3_VP_21mer_3'mf | V(mU)#(fU)#(mA)(fA)(fU)(fC)(mU)(fC)(mU)(fU)(mU)(fA)(mC)(fU)#(mG)#(fA)#(mU)#(mA)#(mU)#(mA)#(fU) |
| | HTT_S_P3_16mer_DIO | (mU)#(mC)#(mA)(fG)(mU)(fA)(mA)(fA)(mG)(fA)(mG)(mA)(mU)(fU)#(mA)#(mA)-DIO |
| HTT Linear | HTT_AS_P3_VP_21mer_3'mf | V(mU)#(fU)#(mA)(fA)(fU)(fC)(mU)(fC)(mU)(fU)(mU)(fA)(mC)(fU)#(mG)#(fA)#(mU)#(mA)#(mU)#(mA)#(fU) |
| | HTT_S_P3_16mer_DIOTEG_5'-3' | (mU)#(mC)#(mA)(fG)(mU)(fA)(mA)(fA)(mG)(fA)(mG)(mA)(mU)(fU)#(mA)#(mA)(TEG)(mU)#(mC)#(mA)(fG)(mU)(fA)(mA)(fA)(mG)(fA)(mG)(mA)(mU)(fU)#(mA)#(mA) |
| Bi-valent 1 | MSH3_1000_P3_as_21mer_3'mf | V(mU)#(fG)#(mC)(fA)(fG)(fU)(mU)(fU)(mC)(fA)(mG)(fU)(mU)(fU)#(mG)#(fC)#(mU)#(mU)#(mC)#(mA)#(fU) |
| | HTT_AS_P3_VP_21mer_3'mf | V(mU)#(fU)#(mA)(fA)(fU)(fC)(mU)(fC)(mU)(fU)(mU)(fA)(mC)(fU)#(mG)#(fA)#(mU)#(mA)#(mU)#(mA)#(fU) |
| | MSH3/HTT_P3_s_16mer_DIOTEG_5'-3' | (mG)#(mC)#(mA)(fA)(mA)(fC)(mU)(fG)(mA)(fA)(mA)(mC)(mU)(fG)#(mC)#(mA)(TEG)(mU)#(mC)#(mA)(fG)(mU)(fA)(mA)(fA)(mG)(fA)(mG)(mA)(mU)(fU)#(mA)#(mA) |
| Bi-valent 2 | MSH3_1000_P3_as_21mer_3'mf | V(mU)#(fG)#(mC)(fA)(fG)(fU)(mU)(fU)(mC)(fA)(mG)(fU)(mU)(fU)#(mG)#(fC)#(mU)#(mU)#(mC)#(mA)#(fU) |
| | HTT_AS_P3_VP_21mer_3'mf | V(mU)#(fU)#(mA)(fA)(fU)(fC)(mU)(fC)(mU)(fU)(mU)(fA)(mC)(fU)#(mG)#(fA)#(mU)#(mA)#(mU)#(mA)#(fU) |
| | HTT/MSH3_S_P3_16mer_DIOTEG_5'-3' | (mU)#(mC)#(mA)(fG)(mU)(fA)(mA)(fA)(mG)(fA)(mG)(mA)(mU)(fU)#(mA)#(mA)(TEG)(mG)#(mC)#(mA)(fA)(mA)(fC)(mU)(fG)(mA)(fA)(mA)(mC)(mU)(fG)#(mC)#(mA) |
| APOE/JAK1 Studies | | |
| | NTC__P3 exNA_AS | V(mU)#(fA)#(mA)(fU)(fC)(fG)(mU)(fA)(mU)(fU)(mU)(fG)(mU)(fC)(mA)(fA)(mU)(mC)(mA)#(mxU)#(fxU) |
| | Dual_NTC_NTC_P3_S | (mU)#(mU)#(mG)(fA)(mC)(fA)(mA)(fA)(mU)(fA)(mC)(mG)(mA)(fU)#(mU)#(mA)(HEG)(mU)#(mU)#(mG)(fA)(mC)(fA)(mA)(fA)(mU)(fA)(mC)(mG)(mA)(fU)#(mU)#(mA) |
| | APOE AS | V(mU)#(fU)#(mG)(fG)(fA)(fU)(mA)(fU)(mG)(fG)(mA)(fU)(mG)(fU)(mU)(fG)(mU)(mU)(mG)#(mxU)#(fxU) |
| | APOE S | (mC)#(mA)#(mA)(fC)(mA)(fU)(mC)(fC)(mA)(fU)(mA)(mU)(mC)(fC)#(mA)#(mA)-DIO |
| | JAK1_P3 exNA_AS | V(mU)#(fC)#(mA)(fG)(fA)(fU)(mU)(fC)(mC)(fU)(mU)(fU)(mG)(fU)(mA)(fC)(mU)(mU)(mC)#(mxU)#(fxU) |
| | Dual_m-APOE_JAK1_P3_S | (mG)#(mU)#(mA)(fC)(mA)(fA)(mA)(fG)(mG)(fA)(mA)(mU)(mC)(fU)#(mG)#(mA)(HEG)(mC)#(mA)#(mA)(fC)(mA)(fU)(mC)(fC)(mA)(fU)(mA)(mU)(mC)(fC)#(mA)#(mA) |

**Supplemental Table 1: Sequences used for in vivo experiments.**
